# Maternal BMI and Placental Transcriptomic Changes: A Meta-Analysis of Gene Expression at the Maternal-Fetal Interface

**DOI:** 10.64898/2026.06.10.731498

**Authors:** Riya Tangri, Timothy R.H. Regnault, Parsia Shooshtari

## Abstract

**Objective:** Maternal body mass index (BMI) is often used as a measure of metabolic status and increased or decreased maternal BMI is associated with a heightened risk of cardiometabolic diseases across generations. The placenta mediates these maternal metabolic cues; however, its genome-wide transcriptional adaptations in response to maternal BMI remain incompletely defined.

**Methods:** To delineate placental genes, pathways, and interaction clusters whose transcript abundance varies with maternal pre-pregnancy BMI through a genome-wide meta-analysis of human placental RNA-sequencing datasets. Placental RNA-seq reads from four publicly available cohorts (n = 146) were mapped to the GRCh38 reference genome and differentially expressed genes were identified. An independent microarray cohort (n = 19) was re-analysed separately to facilitate cross-platform comparison. Functional enrichment employed GO, KEGG, and STRING protein–interaction resources.

**Results:** Meta-analysis of 146 RNA-seq samples identified eight genes with genome-wide significance in placentae from underweight pregnancies including inflammatory signaling gene MAP4K1 and metabolic enzyme PSPH, while overweight and obese categories revealed nominally significant differential expression. KEGG analysis demonstrated significant downregulation of oxidative phosphorylation with increasing maternal BMI, and protein-protein interaction networks revealed inflammatory mediators as central nodes in overweight and obese groups. Independent microarray validation corroborated key findings, including consistent downregulation of oxidative phosphorylation in obesity.

**Conclusion:** Maternal BMI is associated with placental transcriptomic signatures involving inflammatory, metabolic, and hormonal pathways, with consistent downregulation of oxidative phosphorylation across platforms. This genome-wide meta-analysis provides a reproducible catalogue of BMI-responsive placental transcripts that may contribute to developmental programming of offspring health.

**Article highlights:**

- Meta-analysis of 146 RNA-seq samples across four cohorts spanning the BMI spectrum
- Eight genes reached genome-wide significance in underweight placentae
- Oxidative phosphorylation downregulated with increasing maternal BMI across platforms
- Inflammatory mediators emerged as central nodes in obesity-associated PPI networks
- Independent microarray validation corroborated RNA-seq pathway findings

## Introduction

Maternal body mass index (BMI) extremes represent critical modifiers of pregnancy outcomes. Maternal underweight remains prevalent in many populations, affecting approximately 10-20% of pregnancies in some regions [1], while pre-pregnancy overweight and obesity impact an increasingly large proportion of women globally, influencing an estimated 43% of pregnancies worldwide [2] and nearly 25% in North America [3]. Both extremes of maternal BMI are associated with adverse pregnancy outcomes, with underweight linked to preterm birth, intrauterine growth restriction, and low birth weight [1]. Conversely, elevated pre-pregnancy BMI is associated with gestational diabetes and hypertensive disorders [4] and contributes to increased cardiometabolic disease risk in both mother and child across the life course [4, 5].

These intergenerational effects align with the Developmental Origins of Health and Disease (DOHaD) hypothesis, which posits that early-life exposures program lifelong health trajectories through molecular alterations during critical developmental windows. BMI-associated outcomes are thought to be mediated largely via changes in placental development, growth, and function [6, 7]. As the primary interface between maternal and fetal circulations, the placenta interprets and responds to hormonal, nutritional, and inflammatory cues associated with maternal adiposity. Consequently, it must compensate for insufficient maternal reserves in underweight pregnancies while managing excess metabolic substrates in elevated BMI conditions, implying distinct transcriptional programs across the BMI spectrum.

Despite clear evidence that BMI extremes disrupt placental functions, including nutrient sensing, hormonal regulation, cellular energy metabolism and immune surveillance [6, 8], the overall transcriptional changes in the term placenta across the full BMI spectrum remain only partially understood. Bulk-tissue transcriptomic studies, including co-expression network analyses [9], have associated maternal obesity with altered expression of inflammatory, metabolic, and hypoxia-responsive genes [8, 10, 11], with several findings reported to be fetal sex–dependent [6]. However, most available studies arise from individual cohorts that are often underpowered to detect subtle effects and have inconsistent outcomes across datasets [12]. This variability highlights the importance of integrative, cross-cohort approaches capable of distinguishing biological signals from potential study-specific artefacts.

Meta-analytic and data-mining approaches offer a solution to overcoming these issues. Bartho et al. (2021) [13] combined multiple publicly accessible RNA-seq and microarray datasets to highlight mitochondrial regulatory transcripts altered in preterm, post-term, and growth-restricted pregnancies. Likewise, Chabrun et al. (2020) [14] used advanced text-mining and machine learning techniques to integrate placental methylation and transcriptomics data from intrauterine growth restriction (IUGR) pregnancies, connecting coordinated changes in cellular signalling, mitochondrial function, oxidative stress and metabolic pathways. These studies demonstrate that integrating placental omics data through meta-analysis can reveal biologically relevant molecular signatures while overcoming limitations inherent to single-cohort analyses.

Given that metabolic insufficiency and excess likely trigger different compensatory mechanisms in the placenta, a comprehensive analysis across the entire BMI spectrum is essential to understand how maternal metabolic status shapes placental adaptation. Meta-analytic integration across cohorts offers a reliable approach to enhance statistical power, evaluate cross-study reproducibility, and uncover signals hidden by cohort-specific noise [15]. Such tissue-level meta- analyses can contribute significantly to our understanding as they reflect the combined response of all placental cell types and can reveal integrated changes that might be lost when analysing individual cell populations in isolation. To address this gap, we conducted a meta-analysis of four RNA-seq datasets (n = 146) along with an independent microarray cohort (n = 19), covering the full BMI spectrum from underweight to obese. This meta-analysis aimed to establish a reproducible molecular signature of maternal BMI in the placenta, identifying potential biological networks through which maternal metabolic health may influence fetal development. Specifically, we aimed to: (i) identify transcripts whose expression correlates with maternal BMI across the full spectrum, from underweight through obesity; (ii) elucidate enriched biological pathways and co-expression networks; and (iii) compare findings across two technology platforms. This approach will enable identification of both individual gene signatures and systems-level adaptations that may mediate the intergenerational transmission of metabolic disease risk, providing mechanistic insights into the DOHaD paradigm associated with maternal BMI.

## Methods

### Data acquisition

Public repositories (NCBI Gene Expression Omnibus (GEO), Sequence Read Archive (SRA), Database of Genotypes and Phenotypes (dbGaP)) were searched from their inception to May 31st 2023 using the terms placenta, transcriptome, and maternal BMI. Inclusion criteria were: (i) term placentas (≥37 weeks gestation), (ii) available RNA-seq or microarray mRNA data, (iii) recorded maternal pre-pregnancy BMI with supporting metadata. Twenty candidate studies were screened; five met criteria and data-use permission was obtained from primary investigators when data was not publicly available. Exclusion at sample level removed cases lacking phenotypic data or with known confounders (smoking, non-European ancestry, maternal age outside 18-40 years, maternal co-morbidities, pre-term delivery <37 weeks). Final sample sizes are summarised in Table 1.

**Table 1:** Summary of datasets included in the meta-analysis.

| Dataset ID | Platform | n | Underweight | Normal | Overweight | Obese | Reference |
| --- | --- | --- | --- | --- | --- | --- | --- |
| SRP095910 | RNA-seq | 106 | 2 | 61 | 31 | 12 | Deyssenroth et al., 2017 |
| GSE77085 | RNA-seq | 14 | 1 | 9 | 4 | 0 | Buckberry et al., 2017 |
| Suppl. data | RNA-seq | 18 | 1 | 11 | 4 | 2 | Verheecke et al., 2018 |
| GSE56524 | RNA-seq | 10 | 2 | 6 | 1 | 1 | Metsalu et al., 2014 |
| GSE75010 | Microarray | 19 | 0 | 12 | 3 | 4 | Leavey et al., 2017 |

Total RNA-seq n = 146; microarray n = 19. BMI categories defined according to WHO criteria.

### RNA-seq preprocessing

For dataset 1 (Table 1), we obtained dbGaP controlled access and downloaded raw SRA files (dataset 1) in FASTQ format via SRA Toolkit v3.1 [16, 17] using the prefetch and fasterq-dump commands. Read quality was assessed with FastQC v0.11.9 [18]. Adapters were trimmed using Trim Galore Cutadapt tool v0.6.10 [19] with default settings using ASCII+33 quality scores as Phred scores, stringency setting of 1, maximum error rate of 0.1, and a threshold of minimum length 20 bp. Reads were aligned to *GRCh38* (GENCODE v43) with Rsubread v2.14.2 [20] and summarised at gene level with feature Counts function of Rsubread package. Publicly available count matrices (datasets 2–4) were converted from GRCh37 to GRCh38 coordinates using UCSC liftOver [21].

Genes with <10 counts in ≤2 samples were discarded to remove lowly expressed genes that could introduce noise. Libraries with <10,000 total counts and samples with outlier library sizes (<1.5 of interquartile range (IQR) from the first quartile or >1.5 of IQR from the third quartile) were excluded as potential quality outliers. All four datasets were merged on intersecting genes and batch effects were corrected with ComBat (sva v3.46.0) [22]. As RNA sequencing and microarray gene expression platforms are different, with the former sequencing the whole transcriptome while the latter profiles a predefined set of transcripts through hybridization, only the RNA-seq datasets were pooled together.

### Microarray preprocessing

The microarray dataset (dataset #5) was re-analyzed separately to discover other potential significant genes and pathways and to compare the results between the two transcriptomic platforms. Microarray data downloaded from NCBI was already processed, normalized, and converted into log2 values using Affy library from Bioconductor [23].

### Univariate differential expression analysis

#### RNA seq

Differential expression analysis was performed using DESeq2 in R [24]. A negative binomial generalized linear model assessed gene expression changes by pre-pregnancy BMI, modeled both continuously (linear) and categorically (non-linear). BMI categories were: underweight (<18.5), normal (18.5–<25; reference category), overweight (25–<30), and obese (≥30) [25]. Models were adjusted for maternal age, delivery method, and infant sex. Wald statistics were adjusted via Benjamini–Hochberg (BH) procedure; false discovery rate (FDR) <0.05 deemed significant. For exploratory analyses, nominal p-value <0.01 was used.

#### Microarray

Differential expression analysis was conducted using limma v3.58.1 [26] applying an empirical Bayes–moderated linear modeling approach, with identical covariates and significance thresholds.

### Gene-set enrichment and protein-protein interaction

Significant genes (nominal p < 0.01) were analysed with g:Profiler [27] for Gene Ontology (GO) and Kyoto Encyclopedia of Genes and Genomes (KEGG) pathway enrichment. Terms with g:SCS (Set Counts and Sizes) adjusted p <0.05 were reported. Protein–protein interaction networks were generated in STRING v12.0 [28] using a minimum confidence threshold of 0.4, with interactions derived from text mining, experiments, databases, co-expression, neighbourhood, gene fusion, and co-occurrence.

## Data availability

Processed data and analysis scripts are available at [https://github.com/shooshtarilab/Maternal-BMI---Placental-Transcriptome]. Raw sequencing data (datasets #1-4) are available through their original accession numbers listed in Table 1.

## Ethics statement

All data were derived from previously published studies with institutional ethics approval and deposited in public repositories.

## Results

### Participant Characteristics

The meta-analysis comprised 146 RNA-seq participants following quality control and harmonization across datasets. The cohort captured the full BMI spectrum with distinct clinical characteristics across groups (Table 2). For the microarray dataset, a total of 19 participants were included in this current study. The women had a mean BMI of 24.0 kg/m^2^ (range 17.0-42.7) and mean age of 29.1 years (18-40). The placentae were collected from 9 male infants (47.4%) and 6 (31%) of the pregnancies had vaginal delivery.

**Table 2:**
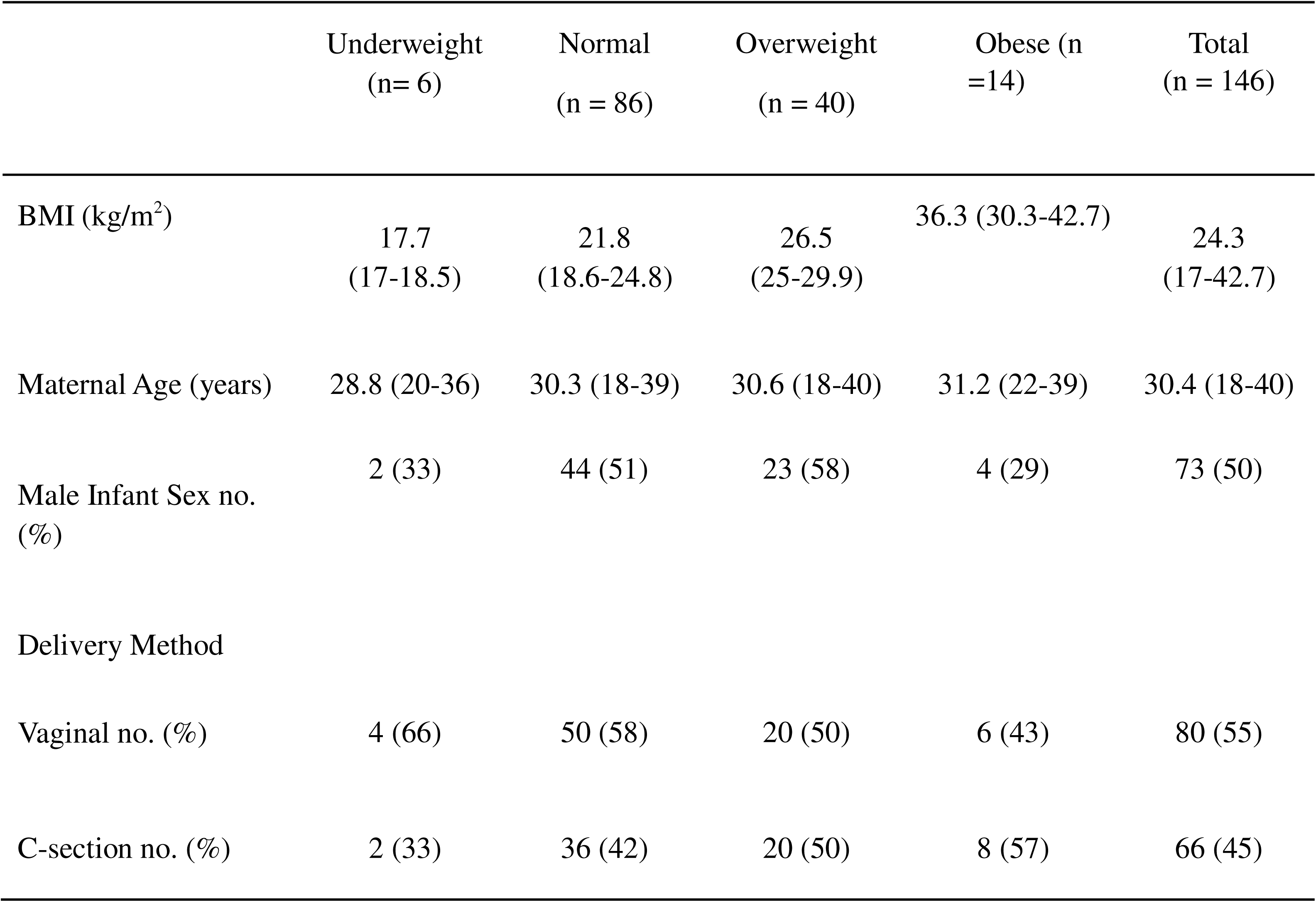
Maternal characteristics of women whose placental samples were used for mRNA analyses.

### Differential Gene Expression Analysis of Pooled RNA-seq Data

Quality control yielded 11,068 overlapping genes across the four RNA-seq datasets for differential expression analysis. Comparing placental transcriptomes across BMI categories revealed eight genes with genome-wide significance (BH-adjusted p < 0.05) in the underweight group: AP3B2, CPXM2, DONSON, GAP43, PARD6G, PSPH, TUBD1, and MAP4K1. While neither overweight nor obese categories or linear BMI analysis yielded genes passing multiple-testing correction, the use of unadjusted p-values revealed nominally significant (p-value < 0.01) genes differentially expressed associated with changing maternal pre-pregnancy BMI (Figure 1).

**Figure 1:**
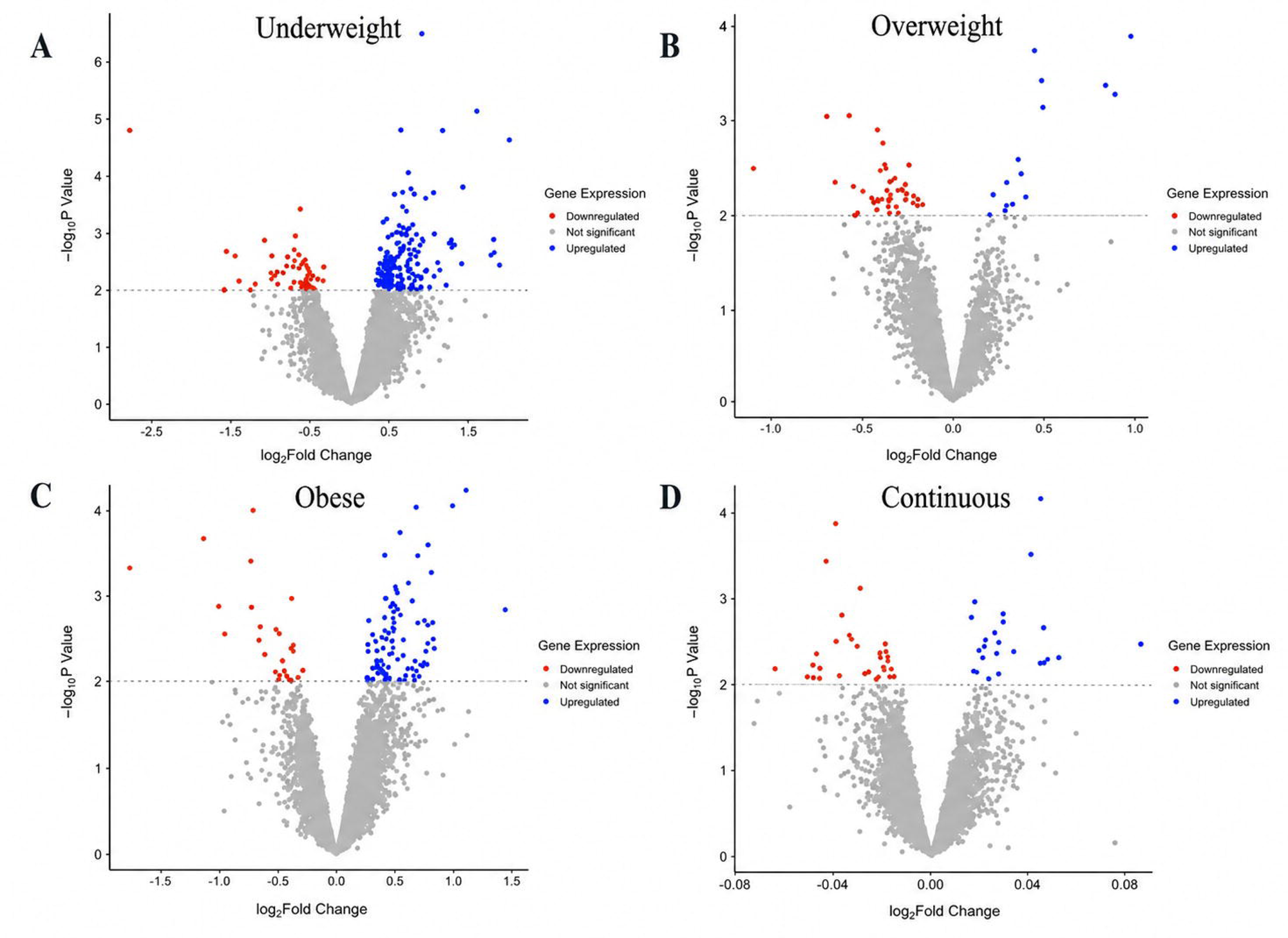
Volcano plots of differential gene expression analysis. Log2-fold changes vs -log10(p values) in mRNA expression associated with (A) maternal pre-pregnancy underweight status; (B) maternal pre-pregnancy overweight status; (C) maternal pre-pregnancy obese status; and (D) continuous maternal pre-pregnancy BMI variable. Data points coloured in red indicate genes nominally significant (p-value < 0.01) and up-regulated. The blue data are nominally significant and down-regulated. Genes not significantly associated with the BMI status are coloured in grey.

When compared to the normal weight status, a total of 221 genes were nominally significant for underweight status (24% genes were downregulated), 58 for overweight (74% downregulated), and 123 for obesity (20% downregulated). Linear analysis identified 55 differentially expressed genes (56% downregulated). Multiple genes showed alterations across conditions (Table 3), with complete results in Supplementary Data 1.

**Table 3:**
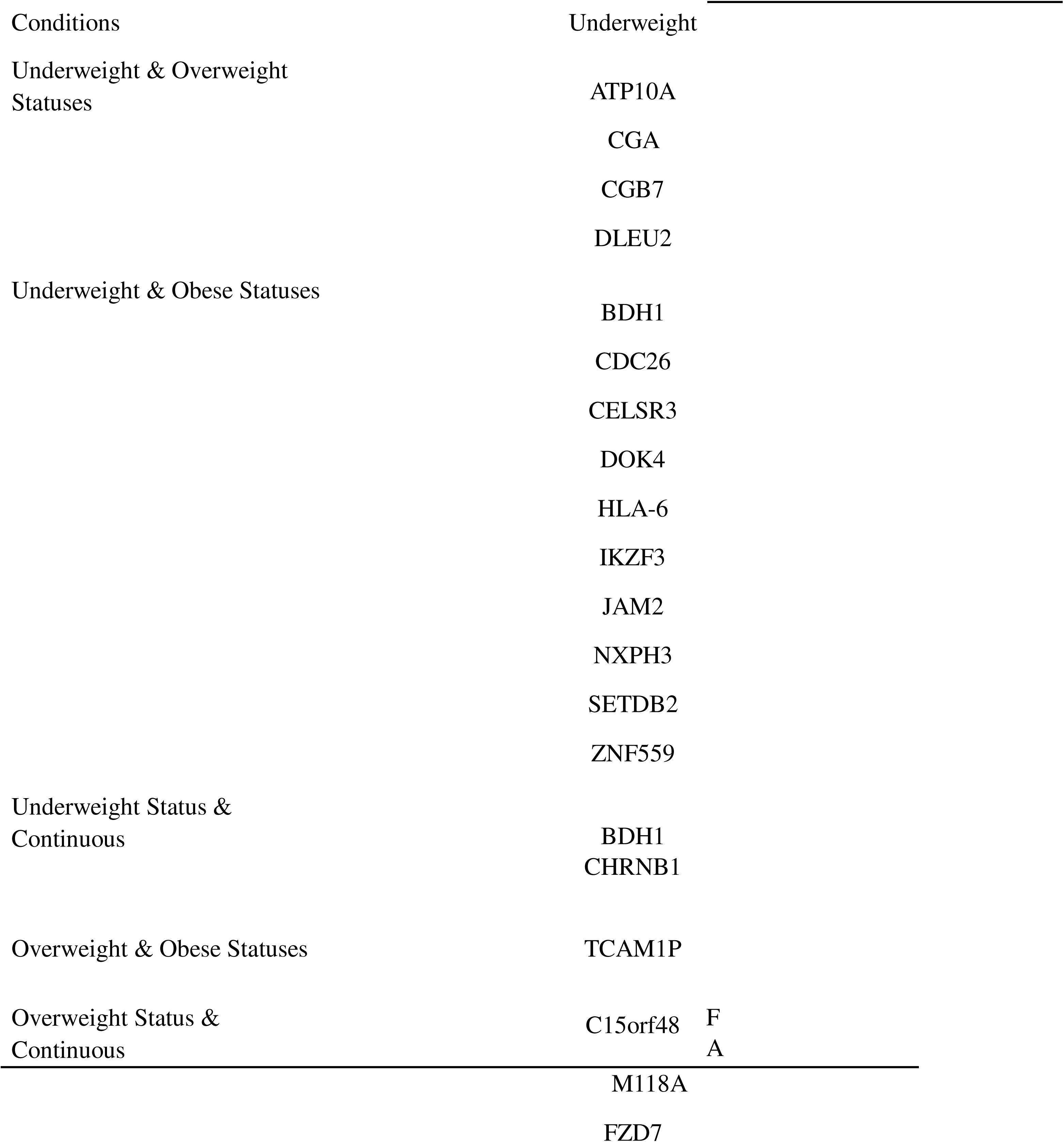
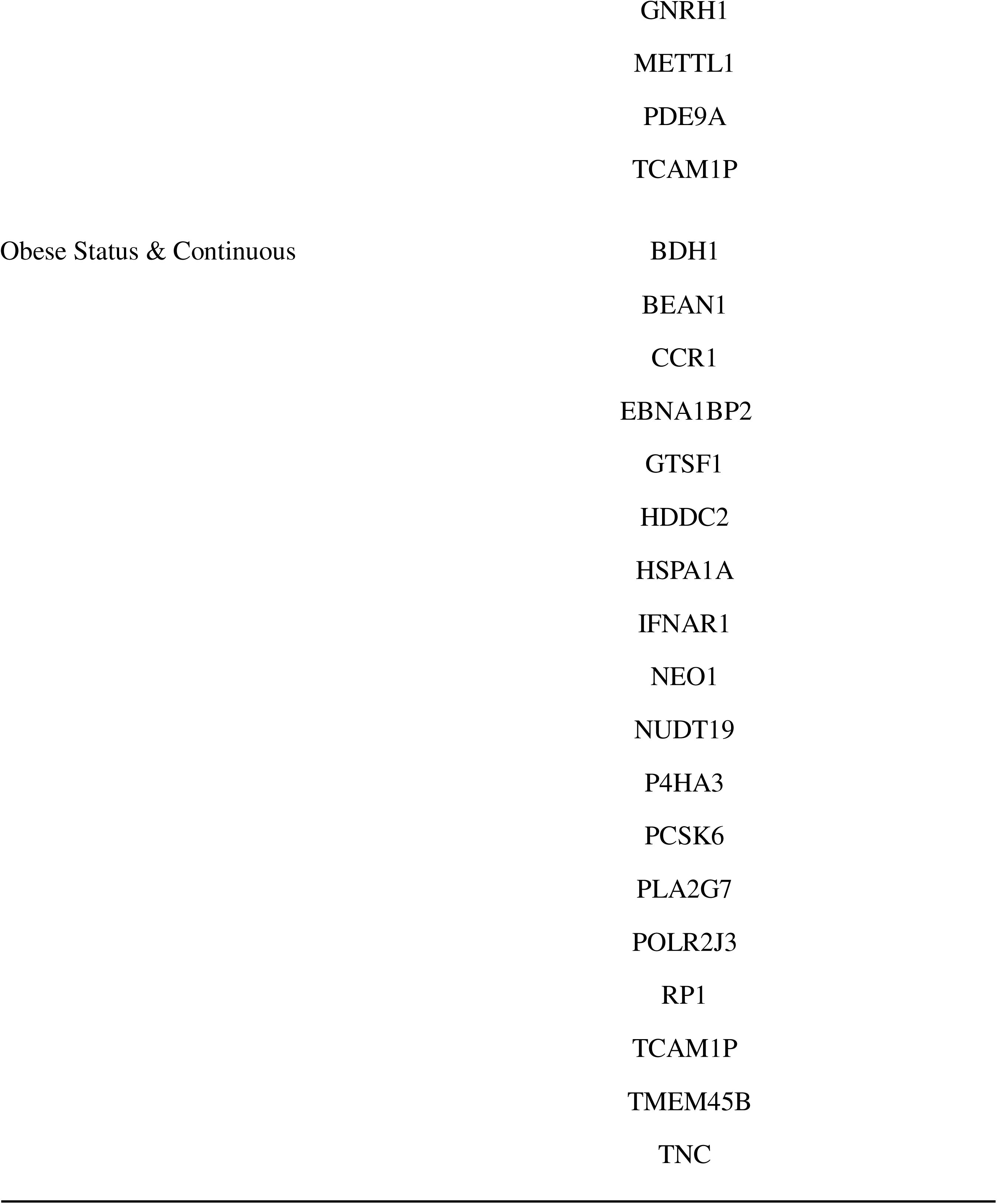
Nominally significant genes differentially expressed in more than one condition.

### GO Pathway Analysis of RNA-seq Data

Pathway enrichment analysis identified biological processes affected by BMI-associated transcriptional changes. While no pathways survived multiple-testing correction, nominal significance (p < 0.01) revealed 77, 237, and 108 enriched GO terms for underweight, overweight, and obese categories, respectively. The ten most significant pathways per category (p < 0.001) are visualized in Figure 2, highlighting key biological processes including hormonal, immune, and metabolic activity, that are affected by maternal BMI

**Figure 2:**
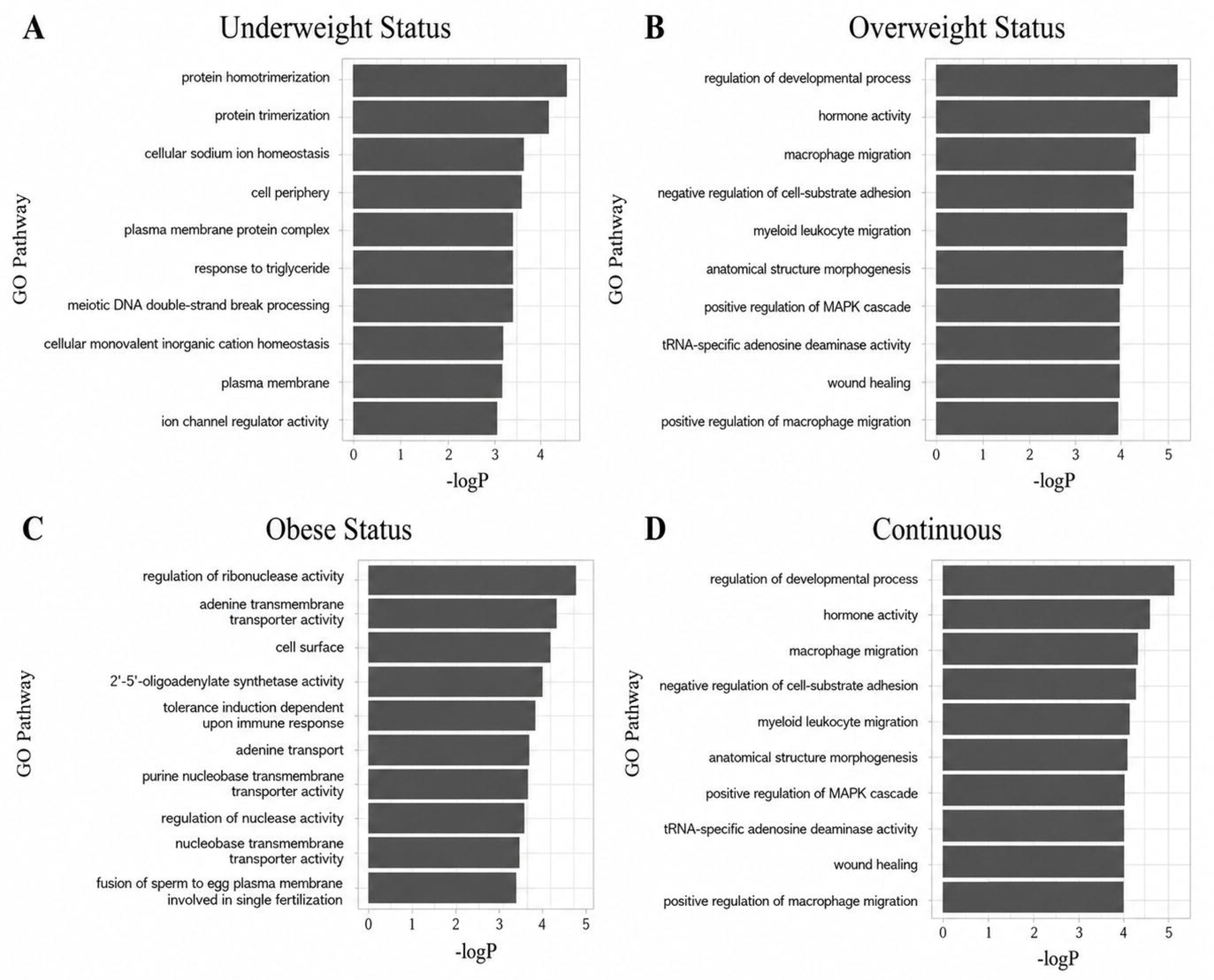
Top 10 most significant GO pathways. Significant GO pathways associated with (A) maternal pre-pregnancy underweight status; (B) maternal pre-pregnancy overweight status; (C) maternal pre-pregnancy obese status; and (D) continuous maternal pre-pregnancy BMI variable.

### KEGG Pathways Analysis of RNA-seq Data

Systematic functional analysis via KEGG identified significantly downregulated pathways (FDR < 5%) across BMI categories (Figure 3), in particular oxidative phosphorylation as maternal BMI increases. No pathways showed significant upregulation.

**Figure 3:**
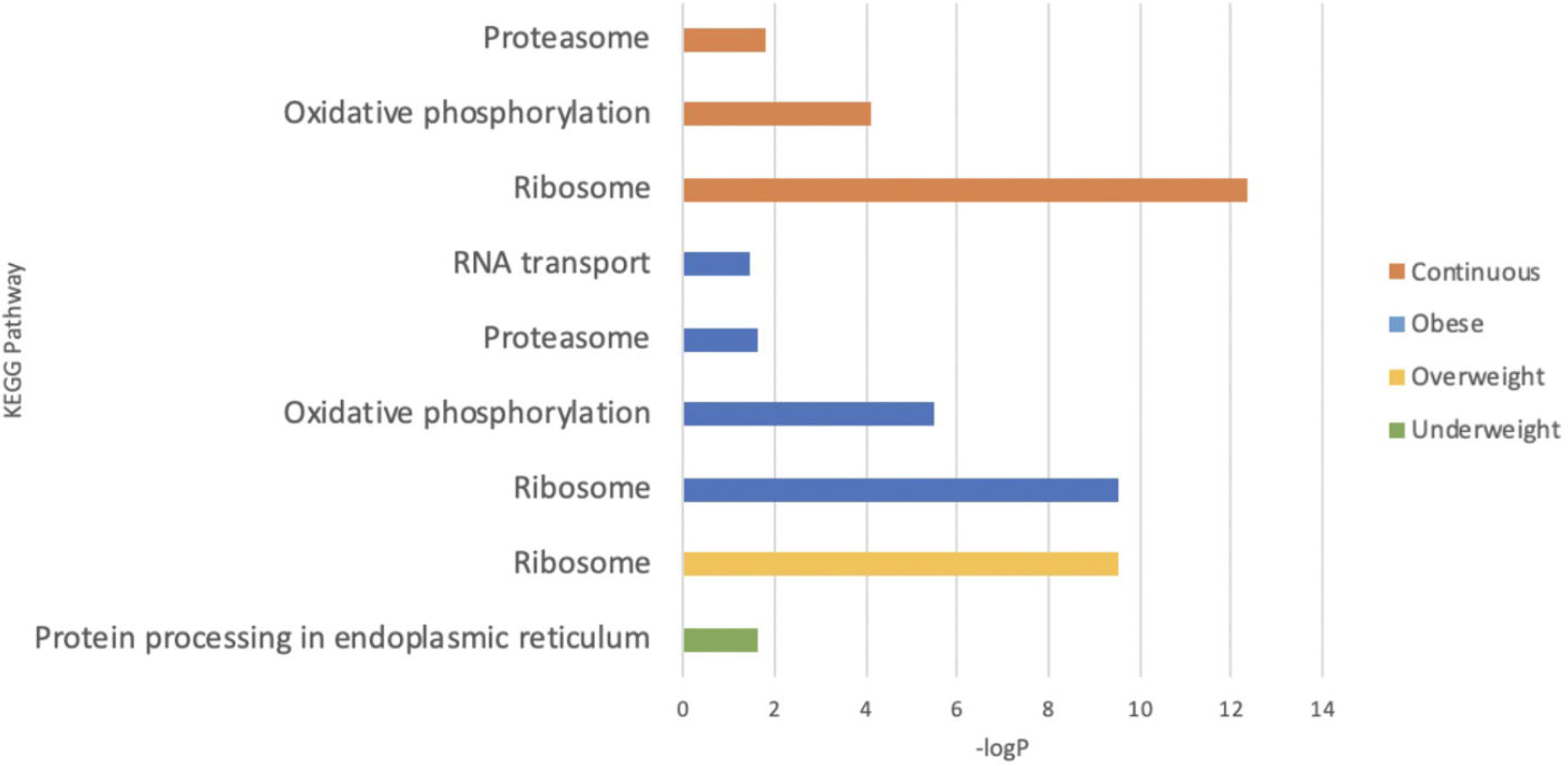
Significant KEGG down-regulated pathways. Orange pathways are associated with the differentially expressed genes associated with the continuous maternal pre-pregnancy BMI variable, blue with maternal pre-pregnancy obese status, yellow with maternal pre-pregnancy overweight status, and green with maternal pre-pregnancy underweight status.

### Analysis of PPI Networks

STRING analysis revealed differential network connectivity across BMI categories. Underweight and continuous BMI analyses showed no significant functional enrichment. However, overweight status generated a highly interconnected network (p < 0.001) with eight functional modules, while obesity produced eleven significant modules (PPI enrichment p < 0.001; Figure 4), notably involving inflammatory mediators such as CCR1 and PLA2G7. Key functional modules in the overweight network included angiogenesis, hormone activity, and extracellular matrix organization, while obesity-associated modules encompassed interferon signaling, immunoglobulin-related pathways, and glycoprotein networks (Figure 4).

**Figure 4:**
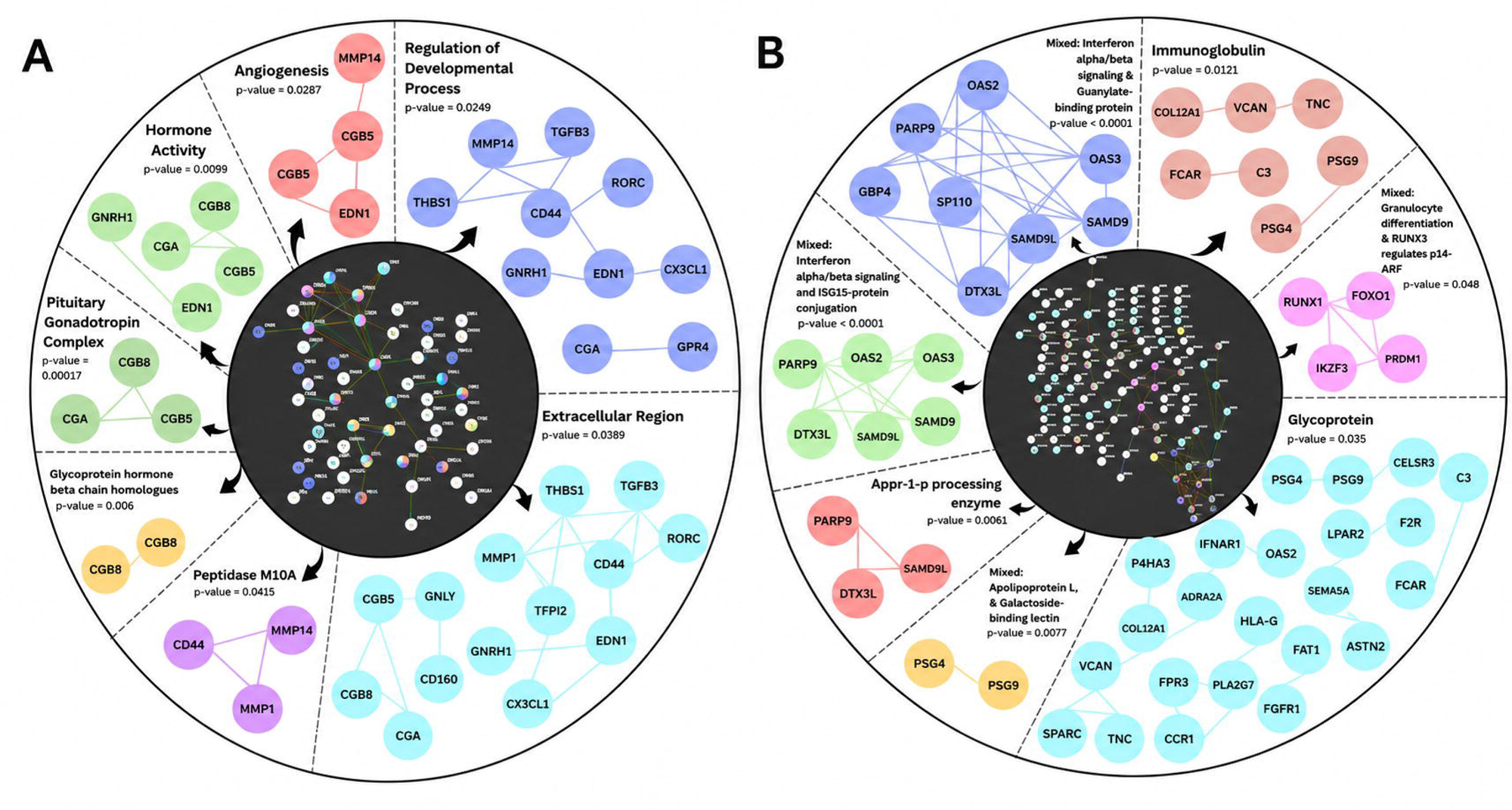
The inner circle represents all the proteins provided as the input and the outer circle presents the significant PPI networks associated with (A) the overweight status; and (B) the obese status, with their respective PPI enrichment p-value. Each node presents a protein and each line represents an interaction. If similar networks in terms of proteins interacting and overall function were determined to be significant, the pathway with the highest p-value was displayed. A complete list of pathways can be found in Supplementary Data 2.

### Analysis of Microarray Data

Independent validation using microarray technology (n = 19) corroborated key findings. Following preprocessing, 20,142 genes underwent differential expression analysis. Using identical significance thresholds (p < 0.01), we identified 58 differentially expressed genes for overweight (31% downregulated), 87 for obesity (28% downregulated), and 98 for continuous BMI (33% downregulated). (Supplementary Data 1). Cross-platform validation revealed six overlapping genes shared between RNA-sequencing and microarray analyses (B3GALT5, CGA, F2R, HBG2, IER3, and NR2F2), originating from the same analytical category in both platforms and exhibited concordant directionality of differential expression.

GO analysis identified 60 nominally significant (p < 0.01) pathway overlaps for the overweight status, 57 pathways for the obese status, and 59 pathways in relation to BMI as a continuum. KEGG analysis revealed two significant pathways (FDR < 0.05): upregulated olfactory transduction in overweight samples and downregulated oxidative phosphorylation in obesity, the latter consistent with RNA-seq findings. PPI network analysis showed no significant enrichment in the microarray dataset.

## Discussion

In this multi-cohort, cross-platform meta-analysis of term placental transcriptomes, we identified a reproducible set of BMI-associated transcriptional signatures involving three major biological domains: mitochondrial energetic, immune/inflammatory signaling and endocrine activity. Although only a small number of transcripts reached genome-wide significance, exclusively within the underweight group, broader nominal changes were observed across overweight, obese, and continuous BMI models. These findings are consistent with prior reports of heterogeneous transcriptional responses that are often subtle and context-dependent to maternal adiposity, and they emphasize the need for large, integrative datasets to detect coordinated biological patterns that may be missed in single-cohort studies.

The strongest FDR-significant signal emerged in the underweight group, in which eight genes showed clear differential expression. No genome-wide significant genes were observed in overweight or obese groups or in continuous modeling, likely reflecting limited statistical power after multiple-testing correction. Prior work investigating the molecular link between maternal BMI and gestational outcomes have produced similar results and resorted to utilizing unadjusted p-values. Clark et al. [11] identified a significant association between maternal pre-pregnancy BMI and placental transcripts for only underweight women, while a study by Cox et al. [9] produced no statistically significant genes. Nevertheless, nominally significant genes across BMI mapped to pathways central to placental adaptation, suggesting physiologic relevance even below stringent thresholds. For example, obese placentae exhibited nominal alterations in lipid and inflammatory mediators, including interferon-stimulated genes, while overweight placentae showed enrichment for hormone-related transcripts. These patterns mirror earlier findings that maternal adiposity subtly reshapes the placental transcriptome rather than producing large, uniform shifts.

Among the eight genes reaching genome-wide significance in underweight placentae, MAP4K1 and PSPH warrant particular attention. MAP4K1 (mitogen-activated protein kinase 1) activates JNK signalling cascades that regulate cellular stress responses and inflammation [29], and its upregulation in underweight pregnancies may reflect compensatory inflammatory signalling under conditions of nutrient scarcity. PSPH (phosphoserine phosphatase), the terminal enzyme in L-serine biosynthesis, is essential for one-carbon metabolism and nucleotide synthesis [30]. Altered PSPH expression could affect serine availability for fetal development, as serine serves as a precursor for glycine, cysteine, and sphingolipids that are critical for membrane biogenesis and neural development. CPXM2, a catalytically inactive metallocarboxypeptidase implicated in cell–cell interactions, has previously been identified as upregulated in placentae from growth-restricted pregnancies [31], suggesting a potential role in placental responses to impaired nutrient supply. DONSON, an essential regulator of DNA replication fork stability, is of interest because mutations in this gene cause microcephalic primordial dwarfism with severe intrauterine growth restriction [32], suggesting that replication stress responses may be engaged in underweight pregnancies. The remaining significant genes, including GAP43, PARD6G, AP3B2, and TUBD1, have established roles in neuronal development, cell polarity, vesicular trafficking, and centriole function, respectively [33, 34, 35, 36], but reports on their relationships with BMI status are lacking. Their identification in underweight placentae represents a novel finding that warrants further investigation to determine whether these reflect direct metabolic responses or indirect effects of altered placental development.

Additionally, it is of interest that several transcripts showed altered expression across multiple BMI comparisons (Table 3), suggesting they may represent core BMI-responsive genes rather than condition-specific adaptations. BDH1 (3-hydroxybutyrate dehydrogenase 1), which catalyzes interconversion of ketone bodies, was differentially expressed in underweight, obese, and continuous BMI analyses, implicating ketone metabolism as a convergent placental response to metabolic extremes at both ends of the BMI spectrum [37, 38]. Similarly, TCAM1P appeared across overweight, obese, and continuous analyses. These overlapping signatures may represent stable markers of placental metabolic adaptation, warranting further investigation as potential biomarkers of extreme ends of placental BMI response.

Interestingly, the direction of differential expression varied markedly across BMI categories: 74% of nominally significant genes were downregulated in overweight placentae compared to only 20% in obese placentae. This asymmetry suggests that overweight and obese states may trigger qualitatively different transcriptional programs rather than simply reflecting dose-dependent responses along a continuum, potentially reflecting distinct compensatory mechanisms or threshold effects in placental adaptation.

Across BMI categories, gene-set enrichment pointed to consistent involvement of membrane transport and substrate handling, supporting nutrient transport as a primary mechanism through which maternal metabolic status influences placental physiology. Prior studies have demonstrated that maternal obesity modifies the expression of placental glucose transporters (GLUT isoforms) [39], fatty acid transporters (FATPs) [40], and amino acid transporters [41]. While our analysis did not identify these specific transporters at nominal significance thresholds, the enrichment of membrane transport and ion homeostasis pathways in GO analysis (Figure 2) suggests broader perturbation of transmembrane substrate handling that may encompass these systems. The nominal enrichment of ion-transport and sodium-dependent pathways in underweight pregnancies is also noteworthy. As several essential amino-acid transporters (e.g., system A) rely on sodium gradients for function, an altered sodium homeostasis may contribute to reduced amino-acid availability, a known feature of intrauterine growth restriction [42]. Notably, SLC38A2 showed nominal differential expression in underweight placentae, providing evidence for altered System A transporter expression alongside the broader sodium homeostasis pathway enrichment [43]. While these results must be interpreted cautiously due to nominal significance, they highlight the potential importance of membrane energetics and transmembrane gradients in placental adaptation to maternal undernutrition and warrant targeted mechanistic studies.

Mitochondrial and protein homeostasis pathways emerged as consistent biological signatures across analyses and platforms. KEGG analysis revealed significant downregulation of multiple pathways with increasing BMI, including oxidative phosphorylation, ribosome biogenesis, proteasome function, and RNA transport (Figure 3). The consistency of oxidative phosphorylation downregulation, particularly in obese placentae and corroborated in microarray analysis, suggests a reproducible biological signal. These findings align with earlier reports of impaired mitochondrial efficiency, increased oxidative stress, and altered metabolic flexibility in placentae exposed to maternal adiposity [44]. Additional studies have linked maternal obesity to placental lipotoxicity, wherein excess circulating free fatty acids accumulate in placental tissue and promote mitochondrial oxidative damage [10]. The resulting mitochondrial dysfunction compromises ATP generation and biosynthetic capacity, thereby impairing placental hormone production and nutrient transport processes essential for fetal development and growth. Collectively, these data suggest that downregulation of oxidative phosphorylation in obese placentae reflects impaired mitochondrial function secondary to the lipotoxic environment rather than an adaptive metabolic response.

Placental endocrine activity also showed BMI-associated perturbations. Overweight placentae demonstrated nominal downregulation of genes encoding hCG subunits (CGA, CGB5, CGB7, CGB8) consistent with clinical observations of reduced circulating hCG concentrations in pregnancies affected by maternal obesity [45]. Because hCG plays essential roles in angiogenesis, immune modulation, and early placental development, altered expression may contribute to impaired placental signaling and increased susceptibility to pregnancy complications such as preeclampsia [46]. The nominal identification of DOK4, a gene with roles in insulin and IGF-1 signaling [47], across BMI extremes further suggests that endocrine pathways responsive to maternal metabolic cues may be transcriptionally sensitive even at modest effect sizes. Additional studies are needed to clarify the extent to which such endocrine alterations directly influence fetal growth trajectories or reflect secondary adaptations to metabolic stress.

Inflammatory and immune-related pathways were among the most consistently enriched signatures associated with elevated maternal BMI. Genes predicted to influence MAPK signaling, macrophage migration, and leukocyte activation were nominally altered in overweight and obese placentae, reflecting the well-described inflammatory microenvironment induced by adiposity. Maternal obesity is known to increase placental macrophage infiltration, oxidative stress, and cytokine expression, and these processes can activate MAPK cascades that promote apoptosis and proinflammatory gene transcription [38]. Our analyses also highlighted type I interferon responses, including OAS2 and OAS3, as well as enriched protein-interaction modules containing chemokine-related genes such as CCR1 and PLA2G7. In addition to these inflammatory networks, the PPI analysis of overweight placentae identified significant clustering in angiogenesis (MMP14, THBS1, EDN1), hormone activity (GNRH1, CGA, CGB8), and extracellular matrix remodeling modules (Figure 4A). Obese placentae showed even greater network complexity, with eleven significant modules including interferon alpha/beta signaling, immunoglobulin-related pathways, and glycoprotein networks (Figure 4B). The emergence of pregnancy-specific glycoproteins (PSG4, PSG9) within obesity-associated PPI networks aligns with their reported roles in maternal immune tolerance. However, PSGs did not emerge as cross-platform signals at the individual gene level, suggesting heterogeneity across cohorts or timing of expression. Targeted studies should revisit this axis alongside the interferon-chemokine signature. Together, these findings support a model in which maternal adiposity induces an immune transcriptional program involving interferon, chemokine, and MAPK-regulated pathways.

The independent microarray analysis corroborated key RNA-seq themes, including directional consistency in oxidative phosphorylation pathways and overlapping involvement of hormonal, immune, and metabolic processes. At the individual gene level, six genes (B3GALT5, CGA, F2R, HBG2, IER3, and NR2F2) emerged as cross-platform validated transcripts. Several of these genes have established links to placental adaptation and dysfunction. NR2F2, a nuclear receptor involved in trophoblast differentiation and angiogenesis, has been implicated in placental vascular remodeling and metabolic regulation [48, 49]. IER3 and F2R are associated with inflammatory and stress-response signaling, supporting evidence that maternal obesity promotes a chronic low-grade inflammatory placental environment characterized by altered cytokine activity and immune dysregulation [11]. Additionally, CGA, which encodes the alpha subunit of human chorionic gonadotropin (hCG), is a key regulator of placental endocrine signaling, angiogenesis, and maternal–fetal immune interactions, suggesting that altered CGA expression may reflect disrupted placental adaptation in pregnancies affected by abnormal maternal BMI [46, 50]. Collectively, these findings reinforce the concept that elevated maternal BMI is associated with coordinated transcriptional alterations affecting energy metabolism, inflammatory signaling, and endocrine function within placental tissue. Although effect sizes differ and platform-specific noise is expected, the broad alignment across datasets supports biological relevance of the identified pathways.

This study benefits from the largest pooled analysis of placental RNA-seq data across the BMI spectrum to date, rigorous batch correction, harmonized phenotype definitions, and cross-platform validation. Nevertheless, several limitations warrant consideration. First, inherent heterogeneity across cohorts -including sample handling, sequencing protocols, and demographic composition - may contribute to residual variation despite correction. Second, BMI is an imperfect measure of adiposity and metabolic health; although widely used in epidemiologic research, it may obscure important variability in maternal metabolic phenotype. Third, pre-pregnancy BMI was self-reported, introducing potential misclassification. Fourth, restricting samples to term placentae limits causal inference: observed transcriptional differences may reflect late-stage adaptation rather than etiologic mechanisms. Finally, bulk tissue profiling cannot resolve cell-type-specific responses; differences in trophoblast, endothelial, immune, or stromal composition may influence observed signatures.

In summary, by integrating multiple cohorts and platforms, our meta-analysis maps a reproducible placental gene-expression signature of maternal adiposity, encompassing nutrient transport, mitochondrial function, hormonal activity, and immune signalling, that aligns with the hypothesised role of the placenta as mediator of maternal metabolic cues in fetal development. These findings refine the molecular framework for DOHaD-related risk associated with maternal BMI extremes and highlight pathways for new mechanistic and translational studies.

## Author contributions

**RT:** Methodology, Software, Formal analysis, Investigation, Writing - Original Draft

**TR:** Conceptualization, Writing - Review & Editing, Supervision

**PS:** Methodology, Validation, Writing - Review & Editing, Supervision

## Acknowledgements

The authors acknowledge the Anishinaabek, Haudenosaunee, Lūnaapéewak and Chonnonton Nations, whose traditional territories are where it was produced. PS is supported by the Children’s Health Research Institute and Investigator Award from the Ontario Institute for Cancer Research. This research was enabled in part by support provided by the Digital Research Alliance of Canada (alliancecan.ca) and Compute Ontario. We would like to sincerely thank the following authors for their willingness to share additional data upon request: Jia Chen and colleagues for the Rhode Island Child Health Study (SRA repository SRP095910); Claire T. Roberts and colleagues for the study *“Placental transcriptome co-expression analysis reveals conserved regulatory programs across gestation,”*; Frédéric Amant and colleagues for the study *“Genetic and microscopic assessment of the human chemotherapy-exposed placenta reveals possible pathways contributive to fetal growth restriction”.* We sincerely appreciate their contribution to advancing research in this field.

